# TRAF6-IRF5 kinetics, TRIF, and biophysical factors drive synergistic innate responses to particle-mediated MPLA-CpG co-presentation

**DOI:** 10.1101/2020.07.17.207209

**Authors:** P. Pradhan, R. Toy, N. Jhita, E. L. Blanchard, A. Atalis, B. Pandey, P. J. Santangelo, D. M. Shayakhmetov, K. Roy

## Abstract

Innate immune responses to pathogens are driven by co-presentation of multiple pathogen-associated molecular patterns (PAMPs). PAMPs and PAMP-analogs are also used as immune-adjuvants to enhance vaccine efficacy by activating various Pattern Recognition Receptors (PRRs), like Toll-like receptors (TLRs). Various combinations of PAMP adjuvants can trigger synergistic immune responses, but the underlying molecular mechanisms driving that synergy are poorly understood. Here, we used synthetic particulate carriers co-loaded with MPLA (TLR4-adjuvant) and CpG (TLR9-adjuvant) as pathogen-like particles (PLPs) to dissect the signaling pathways responsible for the integrated, dual-adjuvant immune response. PLP-based co-presentation of MPLA and CpG to mouse bone marrow derived antigen-presenting cells (BM-APCs) elicited synergistic Type-I Interferon (IFN-β) and IL-12p70 responses, which were strongly influenced by the biophysical properties of PLPs. Mechanistically, we found that the adapter protein MyD88 and the Interferon-Regulatory-Factor-5 (IRF-5), but not the canonical factors IRF-3 or IRF-7, were necessary for production of both IFN-β and IL12p70. TRIF signaling was required to elicit the synergistic response; the absence of TRIF abolished synergy. Importantly, both the kinetics and magnitude of downstream TRAF6 and IRF5 signaling (TRIF-TRAF-IRF5 pathway kinetics) drove the observed synergy. These results identify not only the key signaling mechanism that cooperates to generate a combinatorial response to MPLA-CpG dual engagement in BM-APCs, but they also underscore the critical role that signaling kinetics and biophysical presentation plays in integrated responses to combination adjuvants.

## Introduction

Immune adjuvants have been widely used to boost the potency of weakly immunogenic vaccines against infectious diseases and cancers (*1, 2*). Alum-based adjuvants, first licensed in the 1920s, are used clinically to induce broad innate immune responses, but their lack of specificity makes them unsuitable for a broad range of vaccines and raises concerns for long term tolerability and potential side effects (*3*). On the other hand, certain molecules from viruses, bacteria, and other parasites - collectively known as pathogen associated molecular patterns (PAMPs) - engage with pattern recognition receptors (PRRs) in mammalian cells, located on the cell surface, on endosomes, and in the cytoplasm, to trigger highly specific innate immune responses (*4*).

Highly successful vaccines in human history are composed of live-attenuated or inactivated pathogens, which present multiple adjuvants and antigens assembled on a particulate structure to the immune system and generate protective immunity(*1, 5*). For emerging vaccines consisting of recombinant proteins, peptides or nucleic acids, various PAMPs are being investigated as adjuvants. The triggering of Toll-like receptors (TLRs) on cell membranes and endosomes, and RIG-I like receptors and cGAS-STING receptors in the cytoplasm, induces potent immunity (*6–9*). Although pathogens carry multiple PAMPs and antigens within a unified structure, most vaccine research with PAMPs historically has involved investigations of antigens with independent soluble adjuvants. Recently, particulate carriers and combination adjuvants have gained increasing interest, in order to better mimic the composition of pathogens and elicit more effective immune responses (*10–13*). The main caveat in combination adjuvant delivery is that immune adjuvants have different solubility and diffusion characteristics. When delivered as soluble molecules in vivo, their concurrent presentation and efficient intracellular delivery to innate immune cells is difficult to achieve, especially when one combines a hydrophobic adjuvant (e.g. MPLA) with a highly hydrophilic adjuvant (e.g. CpG). In addition, rapidly diffusible adjuvants (like CpG) induce acute systemic toxicity and require a particulate carrier to prevent rapid diffusion, reduce toxicity, and enable targeted delivery to immune cells (*14, 15*). Particle systems solve the challenges of intracellular delivery, reduce systemic toxicity, and facilitate the co-delivery of diverse types of antigens and adjuvants (*16–19*). By strengthening immune responses, particle systems can safely enhance vaccine efficacy or drive potent anti-tumor effects (*20–22*).

While some TLR adjuvants are antagonistic to each other (e.g., TLR3 and TLR7), other TLR adjuvant combinations drive synergistic immune responses (*23–25*). For instance, nanoparticles with an influenza antigen and co-encapsulated TLR4 and TLR7 adjuvants drive strong immune responses that result in exceptional protection against the flu (*26*). The combination of adjuvants activating TLR4 and TLR9, both PRRs expressed on Gram-negative bacteria, also drives potent, synergistic immune response (*27*). Activation of TLR4 on the cell membrane and the endosome drives the production of Type I interferons and proinflammatory cytokines through NF-κB activation (*28*). Similarly, TLR9 activation on the endosome induces proinflammatory cytokine and Type I interferon production (*29, 30*). Our group has demonstrated that pathogen-like particles with two clinically relevant TLR adjuvants (MPLA, a TLR4 adjuvant, and CpG, a TLR9 adjuvant) induce synergistic innate immune responses in bone marrow-derived antigen-presenting cells (BM-APCs) and adaptive immune responses *in vivo* (*31*). When the PLPs with adjuvants are delivered along with surface-bound ovalbumin antigen, an enhanced humoral immune response is observed when compared to PLPs with single adjuvants and antigen. One group has postulated that synergy is a result of increased TLR9 recruitment to the endosome after TLR4 activation on the plasma membrane. However, the precise cellular and molecular mechanism enabling TLR4-TLR9 synergy is unknown (*32*).

Here, we investigated the TLR4 and TLR9 signaling pathways to identify the underlying mechanism driving the synergistic innate immune response to *cis* presentation of MPLA and CpG in particulate carriers on mouse BM-APCs differentiated with GM-CSF. We demonstrate that biophysical properties significantly affect the innate immune response – specifically, high CpG density dual-adjuvant loaded particles increases the magnitude of synergistic IFN-β and IL-12p70 responses to MPLA and CpG. We also show that even though the cytokine response from the MPLA-dose, when used as a single-adjuvant, is minimal, the TLR4 signaling arm is necessary for the dual-adjuvant synergistic response. Specifically, we found that the TRIF adaptor protein is required for any synergistic enhancement of MyD88 (upstream) and IRF5 (downstream)-dependent type I interferon and IL-12p70 cytokine response to MPLA-CpG-PLPs. Furthermore, we show that the kinetics and magnitude of downstream TRAF6 and IRF5 signaling events plays an important role in driving the synergistic cytokine responses. These findings provide the fundamental basis of the integrated response to TLR4-TLR9 dual engagement in DCs and motivates kinetic signaling studies in the evaluation of innate immune crosstalk for various combinatorial adjuvant platforms.

## Results

### Density of CpG presentation, but not particle size, influences synergistic cytokine responses in BM-APCs treated with MPLA-CpG carrying pathogen-like particles (PLPs)

/To present MPLA and CpG adjuvants at the same time, we synthesized pathogen-like nano and microparticles (PLPs). The PLPs are PLGA particles with branched polyethylenimine (PEI) conjugated to the particle surface. We have extensively published on the synthesis, characterization, and delivery of single and multiple adjuvants and antigens on these PLPs, both *in vitro* and *in vivo* (*19, 31, 33, 34*). For the current studies, MPLA adjuvant was first encapsulated into the particle during double-emulsion, solvent-evaporation based synthesis. The PLPs were then covalently modified with a monolayer of branched polyethylenimine (bPEI) to impart a positive surface-charge, which enables electrostatic loading of the negatively charged CpG adjuvants on the particle surface – thereby allowing dual-loading of both MPLA and CpG. We synthesized 2 sizes of PLPs, micro (MP) sized and nano (NP) sized, with either single adjuvants (M for MPLA, C for CpG) or dual adjuvants (MPLA + CpG, or MC) (**Figure 1A and Table S1 and S2**). PLPs loaded with dual-adjuvants at an MPLA to CpG ratio of 1:10 induced a slightly higher synergistic immune response than particles with a MPLA to CpG ratio of 1:1 (**Figure S1A-B**). Therefore, throughout the rest of experiments, we used a 1:10 ratio of MPLA to CpG. It should also be noted that the encapsulated MPLA dose was chosen to induce a minimal cytokine response, so that we could study how baseline concurrent TLR4 signaling synergizes with the stronger TLR9 response.

Furthermore, the NPs were synthesized with two different adjuvant densities, i.e. mass of adjuvant per particle, (high density, henceforth shown in figures and text as “Hi” and low density, shown in figures and text as “Lo”) for both MPLA and CpG, while MPs were synthesized with low density for MPLA and high density of the CpG adjuvant. A high density of adjuvant was about six-fold higher than the low density of adjuvant. We did not have MPs with both high- and low-density combinations for MPLA + CpG (like for NPs) due to several technical reasons as explained in detail in **Table S1**. Nevertheless, these PLP designs allowed us to compare both size effects at the same density, and density effects at the same size. To reiterate, total MPLA and/or CpG doses were always kept constant across all experimental and control groups in all studies.

**Figure 1.**
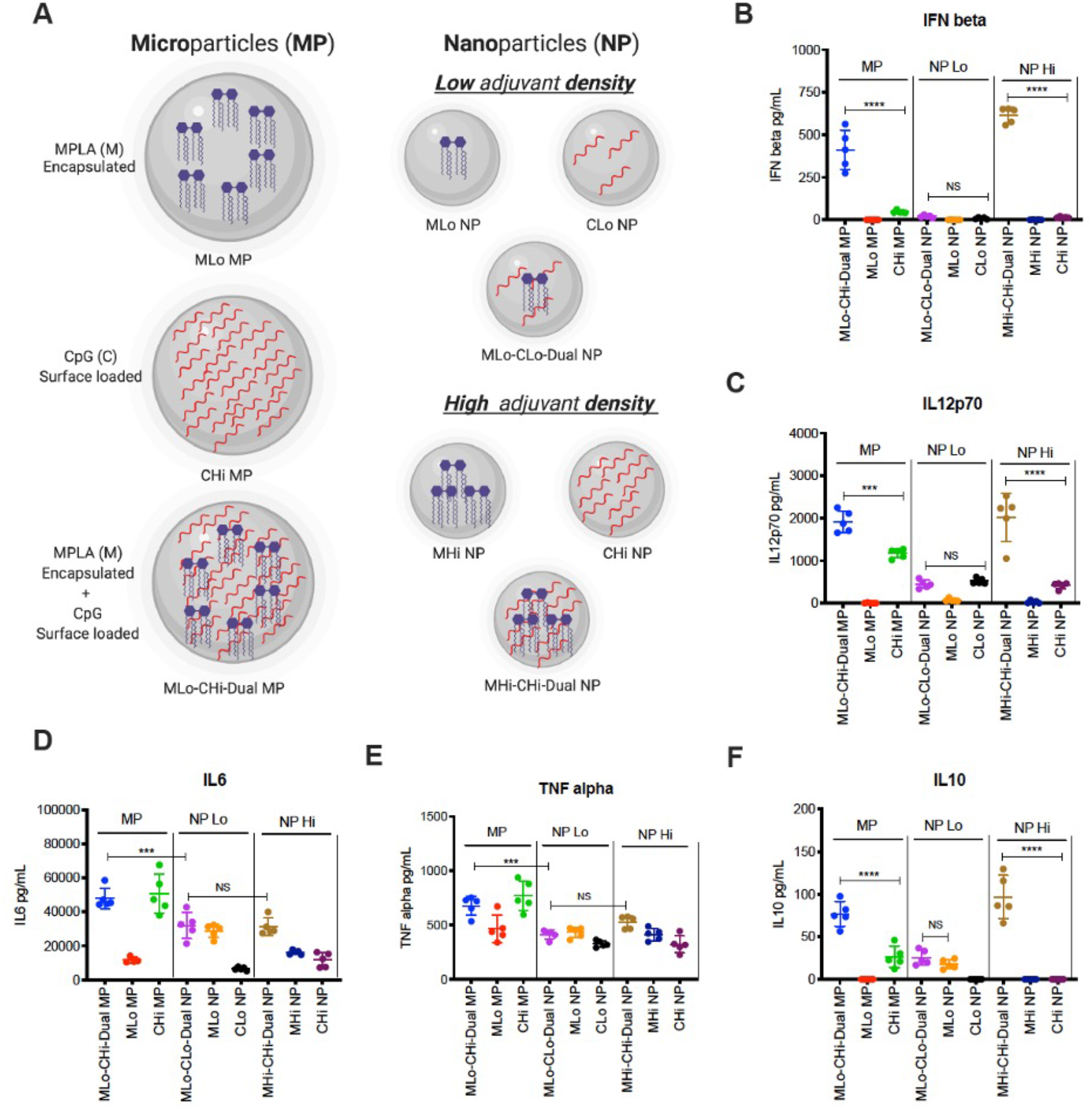
Synergistic cytokine responses from BM-APCs induced by pathogen-like particles (PLPs) with MPLA and CpG depend on CpG adjuvant density. **A)** Schematic of particle formulations for the co-delivery of MPLA and CpG. **B-E)** Murine GM-CSF differentiated murine BM-APCs (300,000 cells/well) were treated with formulations of varying size and CpG ligand density. IFN-β, IL-12p70, IL-6, TNF-α, and IL-10 in cell supernatants 24 hours after treatment. Each data point represents an independently treated well (n=5). Center lines designate the mean value and error bars represent SD. ***P < 0.001, ****P<0.0001, NS-not significant; one-way ANOVA with Tukey’s multiple comparison test.

We first studied how particle size and adjuvant density influenced the cytokine response in GM-CSF differentiated mouse bone marrow-derived antigen presenting cells (BM-APCs). Numerous previously published papers describe GM-CSF differentiated murine bone marrow cells as bone marrow-derived dendritic cells (BMDCs); however, recent literature evidence suggest a heterogenous mixture of innate immune cells, including dendritic cells, macrophages, and monocytes present in the GM-CSF differentiated bone marrow cells around 6-7 days of culture (*35–37*). Hence, we refer our GM-CSF differentiated murine bone marrow cells as BM-APCs instead of BMDCs throughout this manuscript.

In BM-APCs, the IFN-β response to MP (MP-MLo-CHi) and NP-Hi with both MPLA and CpG was synergistic when compared to the IFN-β responses to single adjuvants. In contrast, there were minimal IFN-β responses from NPs with low densities of MPLA and CpG adjuvants (**Figure 1B**). The IL-12p70 responses to MP and NP-Hi with both MPLA and CpG mirrored those of the IFN-β responses. A lower IL-12p70 response was observed after treatment with NPs with a low density of CpG, and concurrent treatment with MPLA did not generate a synergistic response (**Figure 1C**). We also measured IL27, which is one of the members of IL12 cytokine families. As with IL12p70, IL27 response was also synergistically enhanced in BM-APCs by MPLA-CpG-Dual PLPs (both MPs and NPs) with high CpG density (**Figure S2**). Additionally, we measured the IL-6, TNF-α and IL-10 responses to PLPs with MPLA and CpG. IL-6 responses were primarily CpG-driven in MPs, MPLA-driven in NPs with low adjuvant density, and additive between MPLA and CpG in NPs with high adjuvant density (**Figure 1D**). TNF-α secretion was not statistically different between the groups (**Figure 1E**). However, for both IL-6 and TNF-α, MPs showed higher responses compared with NPs (**Figure 1D and E**). As with IFN-β and IL-12p70, the IL-10 response was synergistic for MPs and NPs with high densities of adjuvants, but not synergistic for NPs with low densities of adjuvants (**Figure 1F**).

### The adaptor protein TRIF is required for the synergistic Type I IFN and IL-12p70 response induced by dual MPLA-CpG PLPs in BM-APCs

To identify why PLPs with MPLA and CpG induce synergistic IFN-beta and IL-12p70 responses, we systematically evaluated the signaling pathways driven by TLR4 and TLR9. Activation of TLR4 on the plasma membrane results in the recruitment of the adaptor protein MyD88. When MPLA-CpG-Dual PLPs are internalized, they can activate both TLR4 and TLR9 on the endosome. Endosomal TLR4 activation is known to recruit the adaptor protein TRIF, while endosomal TLR9 activation recruits the adaptor protein MyD88. Both TLR9 and TLR4 signaling, regardless of location, activate TRAF6, which is a central mediator for downstream signaling that ultimately triggers Type I interferon and IL-12p70 (**Figure 2A**).

**Figure 2.**
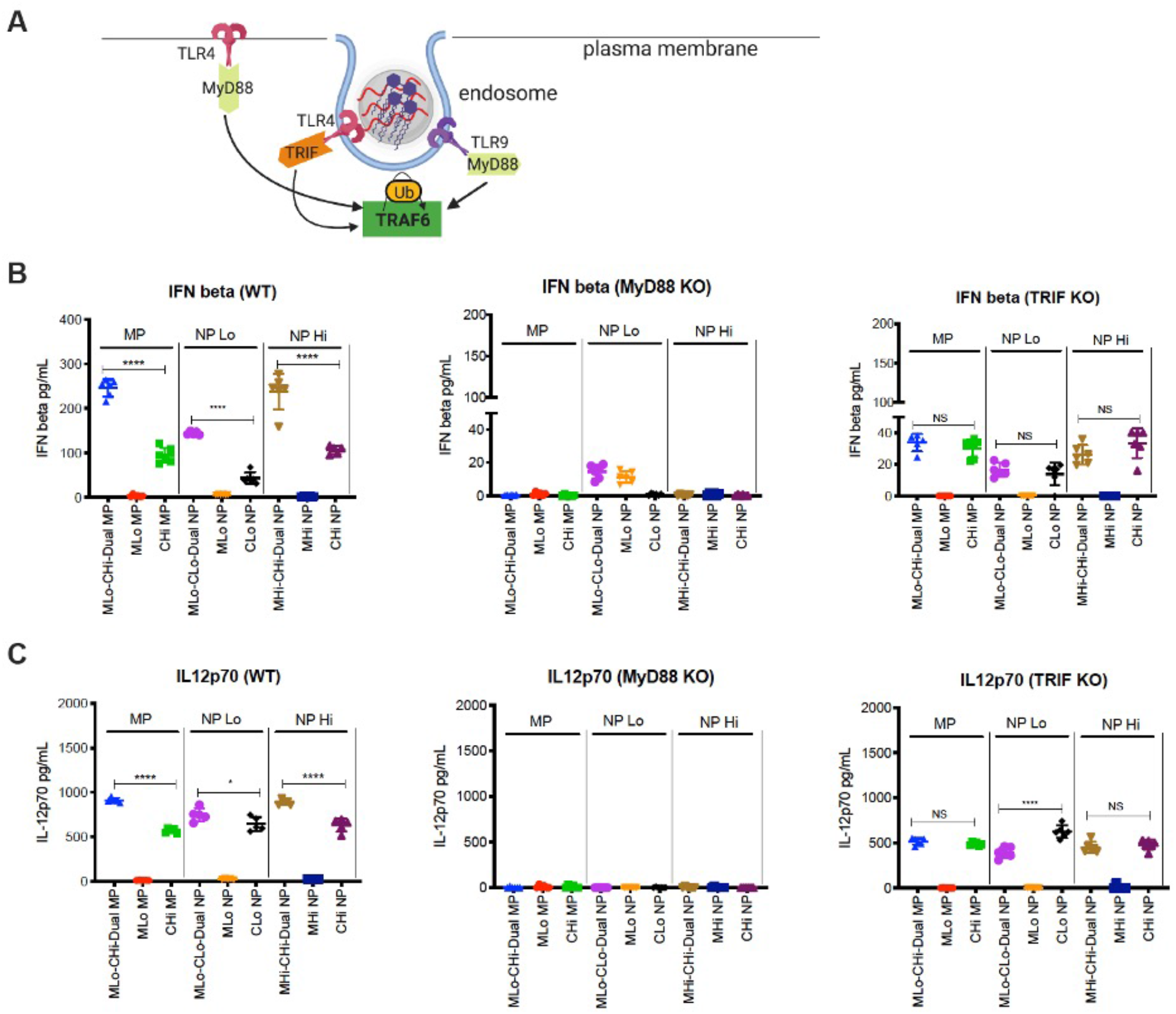
Knockdown of the adaptor proteins TRIF ablates synergy from PLPs with MPLA and CpG in BM-APCs. **A)** Schematic showing early signaling through the adaptor proteins MyD88 and TRIF following activation of TLR4 and TLR9. **B)** IFN-β production from ΒΜDCs derived from wild-type, MyD88^−/−^, and TRIF^−/−^ mice after PLP treatment. **C)** IL-12p70 production from BM-APCs derived from wild-type, MyD88^−/−^, and TRIF^−/−^ mice after PLP treatment. Each data point represents an independently treated well (n=5-6). Center lines designate the mean value and error bars represent SD. *P < 0.05, ****P<0.0001, NS-not significant; one-way ANOVA with Tukey’s multiple comparison test.

Here we first evaluated the effect of MyD88 and TRIF knockdown on the IFN-β and IL-12p70 responses from BM-APCs induced by PLPs. With MyD88^−/−^ BM-APCs, we observed complete ablation of the IFN-β and IL-12p70 response for dual MPLA-CpG-Dual-MPs and NPs with high density for CpG, suggesting that MyD88 is the primary adaptor protein for the MPLA-CpG-dual PLP adjuvant signaling (**Figure 2B and 2C**). In contrast, the synergistic increases in IFN-β and IL-12p70 responses for MPLA-CpG-Dual MPs and NPs with high CpG density were lost in TRIF^−/−^ BM-APCs, and the cytokine levels remained at the CpG-Hi single adjuvant level for MPs and NPs (**Figure 2B and 2C**). This indicates that TRIF adaptor protein is required only for the synergistic enhancement of IFN-β and IL-12p70 responses by dual-loaded MPLA-CpG-PLPs. Furthermore, using BM-APCs from TLR4^mut^ mice, we also confirmed that synergistic IFN-β and IL-12p70 production due to MPLA-CpG-Dual PLPs with high CpG density is dependent on TLR4 signalling from MPLA (**Figure S3A-B**). However, with BM-APCs from TLR9^mut^ mice, we observed synergistic enhancement of IFN-β and IL-12p70 responses (at a lower level than WT BM-APCs) for Dual PLPs with high CpG density (**Figure S3A-B**), which could be due to the lack of complete ablation of TLR9 gene in the TLR9^mut^ mice.

### IRF5 but not IRF3 or IRF7, drives the innate immune response triggered by dual-adjuvant loaded MPLA-CpG PLPs in BM-APCs

Activation of TLR4 and/or TLR9 induces TRAF6 ubiquitination, which proceeds to phosphorylate interferon regulatory factors 3, 5, or 7 (IRF3, 5, or 7, **Figure 3A**). To identify the most important interferon regulatory factor in downstream signaling, we measured IFN-β and IL-12p70 in BM-APCs from IRF3, IRF5 (without DOCK2 gene mutation, **Figure S4**), and IRF7^−/−^ mice. The synergistic IFN-β responses in IRF3^−/−^ BM-APCs were higher than in wild-type cells. In IRF5^−/−^ BM-APCs, IFN-β responses were completely ablated regardless of PLP treatment. There were no differences in IFN-β responses in IRF7^−/−^ BM-APCs when compared to wild-type (**Figure 3B**). As with IFN-β, IL-12p70 responses in IRF3^−/−^BM-APCs were increased when compared to wild-type. IL-12p70 responses were also ablated in IRF5^−/−^ BM-APCs. In IRF7^−/−^ BM-APCs, there were no changes in IL-12p70 secretion when compared to wild-type (**Figure 3C**). We further investigated whether IFN-β increased IL-12p70 secretion by engaging IFN alpha receptor (IFNAR) in an autocrine or paracrine way. In IFNAR^−/−^ BM-APCs, synergistic level of IL-12p70 decreased partially in MPLA-CpG-Dual MP and NP groups, suggesting that the autocrine or paracrine effect of IFN-β have a partial effect on enhancement IL-12p70 production, via IFNAR. Notably, we observed significantly higher IFN-β levels in IFNAR^−/−^ BM-APCs compared to wild-type BM-APCs, likely as a result of accumulation of unused IFN-β in the culture media in BM-APCs from IFNAR^−/−^ mice (**Figure S5A-B**).

**Figure 3.**
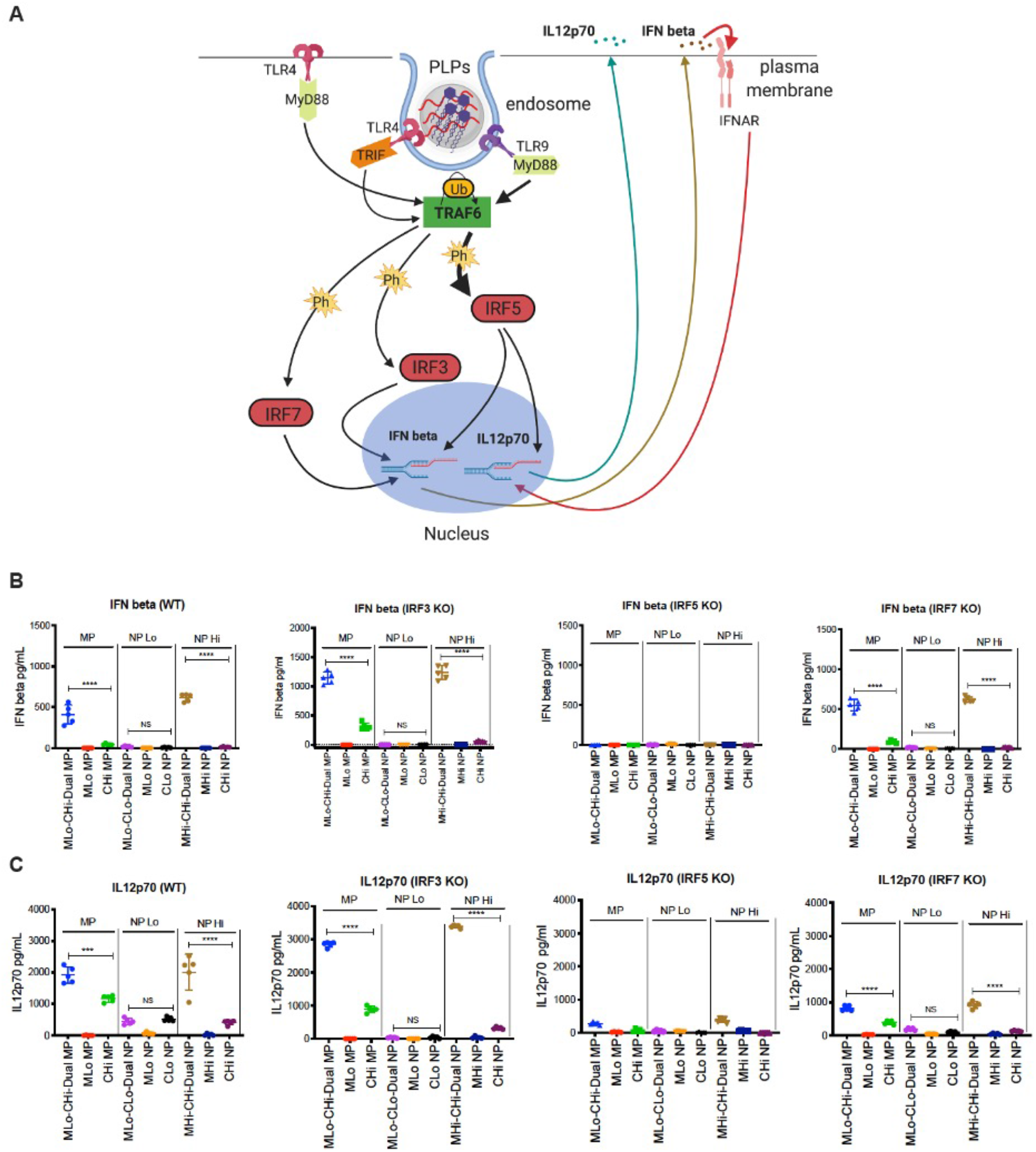
IRF5, not IRF3 or IRF7, drives innate immune response triggered by MPLA and CpG in BM-APCs. **A)** Diagram of TLR4 and TLR9 downstream signaling. **B)** IFN-beta production from BM-APCs derived from WT, IRF3^−/−^, IRF5^−/−^, and IRF7^−/−^ mice. **C)** IL-12p70 production from BM-APCs derived from WT, IRF3^−/−^, IRF5^−/−^, and IRF7^−/−^ mice. Each data point represents an independently treated well (n=5). Center lines designate the mean value and error bars represent SD. ***P < 0.001, ****P<0.0001, NS-not significant; one-way ANOVA with Tukey’s multiple comparison test.

### Sustained and elevated IRF5 phosphorylation and TRAF6 levels are responsible for synergistic innate immune responses by MPLA-CpG-Dual-PLPs in BM-APCs

TRAF6 ubiquitination activates a sequence of kinases, which ultimately leads to IRF5 phosphorylation. This event enables activated IRF5 to translocate from the cytoplasm into the nucleus, where it can initiate transcription of mRNA encoding for IFN beta and the pro-inflammatory cytokine IL-12p70 (**Figure 4A**). Given that we found that IRF5 is primarily responsible for the dual adjuvant signaling, we further investigated what aspect of IRF signaling, amplitude or kinetics or both, is responsible for the synergistic cytokine production. To evaluate this, we measured total IRF5 and phosphorylated IRF5 protein levels over time. After MPLA-MP (single adjuvant) treatment, phosphorylated IRF5 levels were highest after 30 minutes to 1 hour. No phosphorylated IRF5 was detected after CpG-MP treatment. Interestingly, phosphorylated IRF5 levels were high between 30 minutes to 4 hours after MPLA-CpG-Dual-MP treatment (**Figure 4B**), indicating prolonged signaling. In agreement with phosphorylated IRF5 levels, the rate of IRF5 translocation to the nucleus peaked 4 hours after treatment (**Figure 4C**). The ratio of nuclear to cytoplasmic IRF5 in BM-APCs treated with MPLA-CpG-Dual-MP is approximately 5 times higher than the ratio in BM-APCs treated with CpG-MP (**Figure 4C**). A second surge of IRF5 translocation from the cytoplasm to the nucleus occurs at 24 hours. We also assessed the kinetics of IFN-β and IL-12p70 production by BM-APCs of PLP adjuvants and observed synergistic increase in the cytokine production for MPLA-CpG-Dual-MP, which peaked at around 6 hours and sustained over 24 hrs after stimulation with MPLA-CpG-Dual-MPs **(Figure S6).**

**Figure 4.**
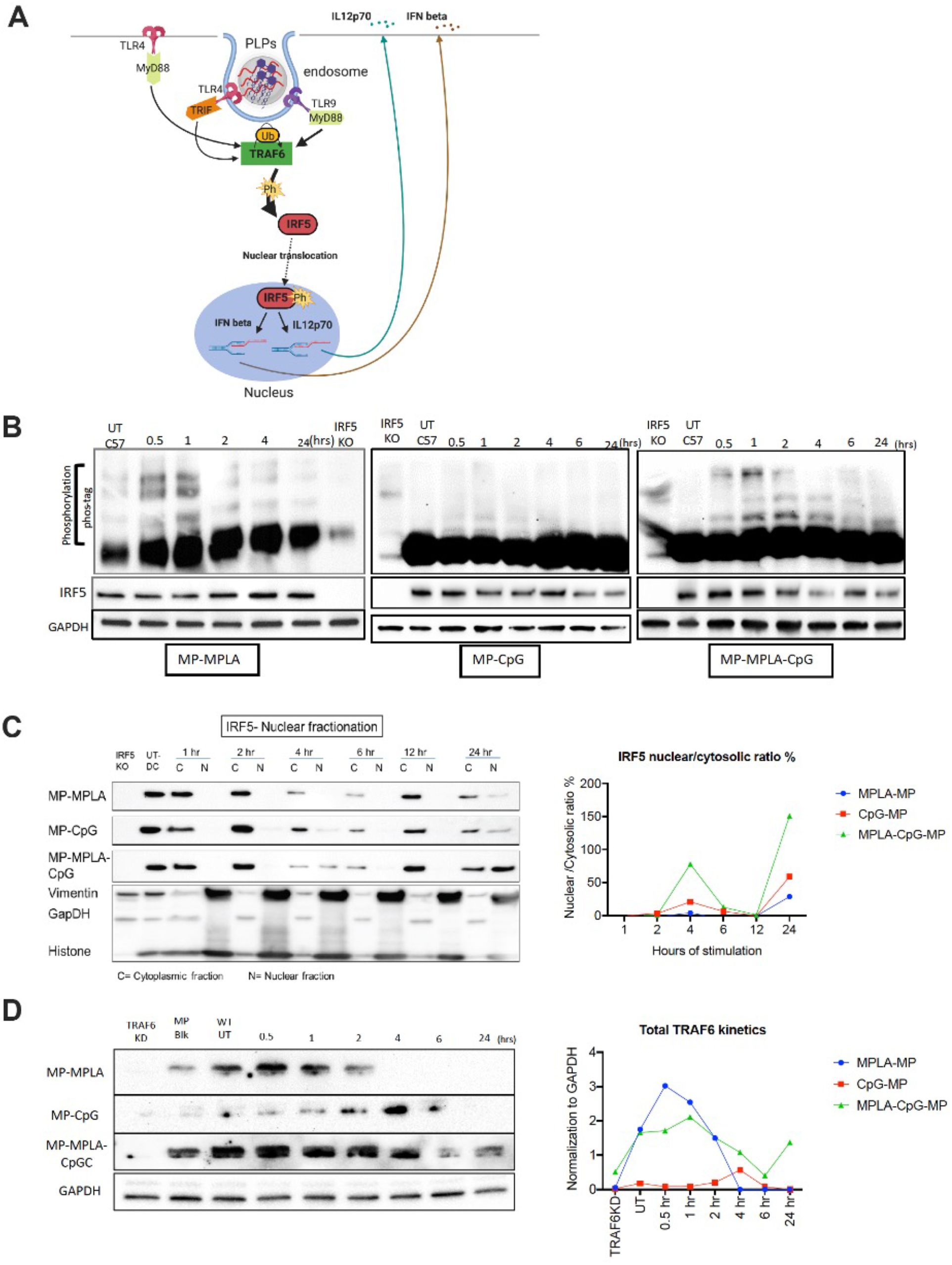
Sustained IRF5 phosphorylation, higher IRF5 nuclear translocation and elevated TRAF6 is responsible for synergistic IFN and IL-12p70 responses in BM-APCs. **A)** Schematic showing TRAF6 and IRF5 signaling following TLR4 and TLR9 activation. **B)** Levels of phosphorylated (Phos-tag) and total IRF5 (SDS PAGE) after 0.5, 1, 2, 4, 6, and 24 hours of BM-APC’s treated with MPs with MPLA and/or CpG. GAPDH was used as a loading control and treated IRF5^−/−^ BM-APCs were used as negative controls. **C)** Total IRF5 levels in the nuclear and the cytoplasmic fractions at 1, 2, 4, 6, 12 and 24 hours of BM-APC treated with MPs with MPLA and/or CpG. Ratio of nuclear to cytoplasmic IRF5 after 1, 2, 4, 6, 12, and 24 hours of treatment, where a higher ratio indicates a higher rate of nuclear translocation. Ratios were performed by Bio Rad Image Lab software. **D)** Levels of total TRAF6 (SDS-PAGE) after 0.5, 1, 2, 4, 6, and 24 hours of BM-APC’s treated with MPs with MPLA and/or CpG. GAPDH was used as a loading control and TRAF6 knocked-down in BM-APCs were used as negative controls. On the right is the graphical representation of the GAPDH normalization of each blot. This was done with Bio Rad Image Lab software.

Next, we measured the expression kinetics of TRAF6, which is the master regulator that drives IRF5, NF-κΒ, and AP-1 signaling and ultimately leads to production of Type I interferon and pro-inflammatory cytokines, including IL-12p70. We hypothesized that sustained and elevated TRAF6 signaling upstream to IRF5 could lead to synergistic IFN-β and IL-12p70 responses in BM-APCs activated by MPLA-CpG-Dual-MP (**Figure 4D**). In BM-APCs treated with MPLA-MP, total TRAF6 expression peaked at 30 minutes after treatment and disappeared after 4 hours. For BM-APCs treated with CpG-MP, total TRAF6 expression peaked at 4 hours and declined by 6 hours. Interestingly, for BM-APCs treated MPLA-CpG-Dual MP, sustained and elevated TRAF6 expression was observed over 24 hrs (**Figure 4D**), which indicates that kinetics of TRAF6 signaling is correlated with cytokine synergy.

Overall, as shown in **Figure 5**, we identified the key signalling mediators that play critical roles in driving synergistic IFN-β and IL-12p70 cytokine responses induced by co-presentation of the TLR4 adjuvant MPLA, and the TLR9 adjuvant CpG on synthetic pathogen-like particles. MPLA-CpG-Dual PLPs activate both TLR4 and TLR9 in the endosome which subsequently engage TRIF (primarily from MPLA) and MyD88 (primarily from CpG) adaptor and lead to sustained and elevated TRAF6 expression and IRF5 phosphorylation and nuclear translocation. Finally, sustained and elevated level of IRF5 causes synergistic production of IFN-β and IL-12p70. Secreted IFN-beta binds to IFNA receptor and further boosts IL-12p70 production by autocrine/paracrine signaling.

**Figure 5.**
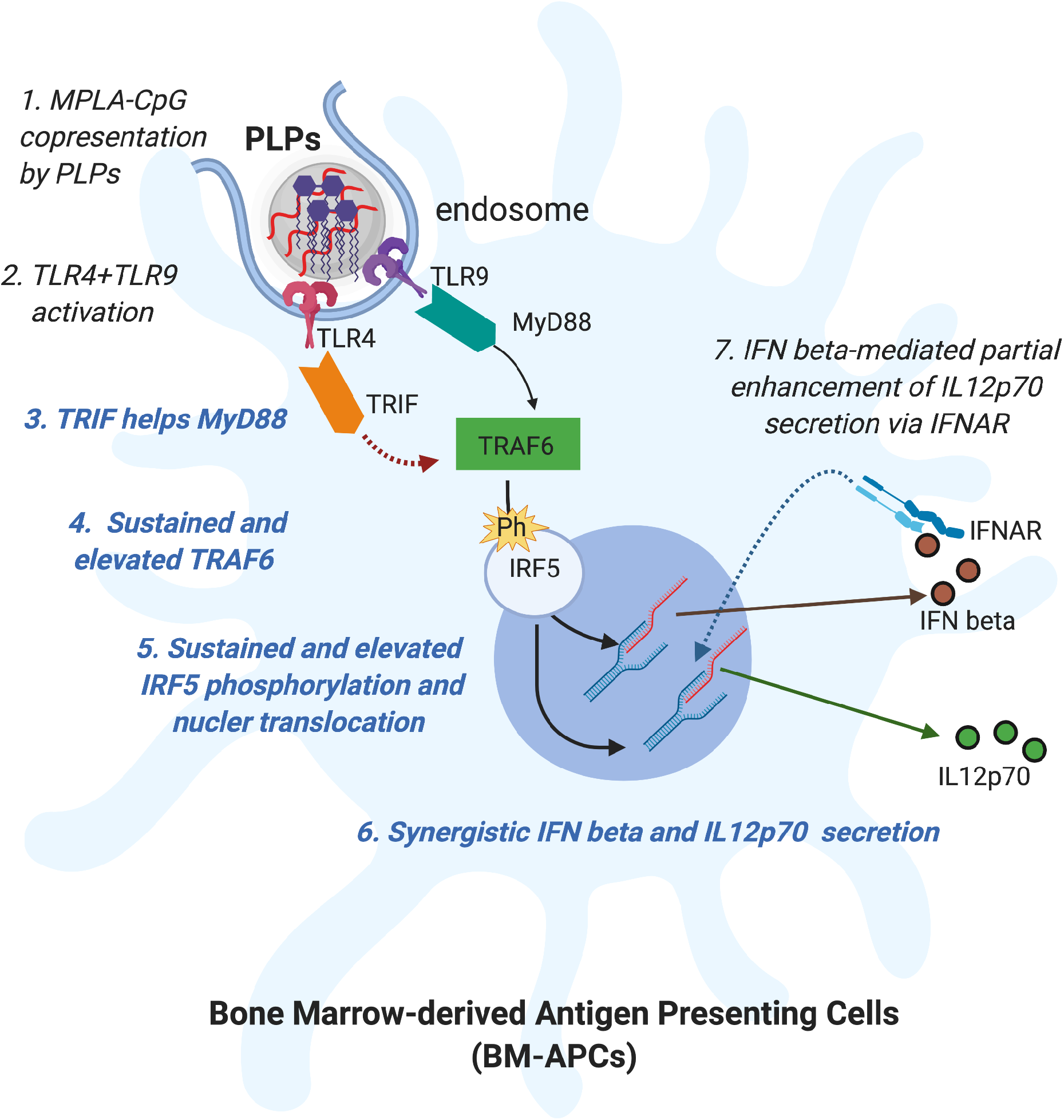
Schematic showing the mechanism of synergistic IFN beta and IL-12p70 cytokine response due to MPLA and CpG co-presentation on PLPs in BM-APCs.

## Discussion

We investigated the effect of biophysical properties such as size and adjuvant density on the efficiency of synthetic particulate carriers (PLPs) to induce synergistic innate immune responses through concurrent activation of TLR4 and TLR9 in BM-APCs. In addition, we investigated how specific intracellular signaling pathways drive synergistic innate immune responses through TLR4 and TLR9 crosstalk. This synergistic phenomenon has been identified in multiple reports by our group and others (*7, 21, 31, 38*). Other have reported that in neutrophils, MPLA and CpG dual-delivery may trigger TLR9 translocation from the cytosol to the endosome by the initial TLR4 activation(*32*). Despite evidence of such synergy, the underlying signaling mechanism and the specific signaling molecules involved in driving MPLA-CpG (TLR4/TLR9) dual-adjuvant synergy in antigen-presenting cells have not yet been fully elucidated. Here we evaluated each of the intermediate steps in the TLR4 and TLR9 signaling pathways in BM-derived APCs to identify if and how they contribute to synergistic Type-I Interferon and IL-12 responses.

We have previously shown that increasing surface density of CpG (as a single adjuvant) on PLPs increased IL-12p70 production from BM-APCs (*33*). Here, most importantly, we found that the adjuvant density effect also applies to IFN-β and IL-12p70 responses from dual TLR4/TLR9 engagement. NPs with a high MPLA and CpG density triggered synergistic immune responses in BM-APCs, while NPs with low MPLA and CpG density triggered a low IL-12p70 response and no detectable IFN-β response in BM-APCs. The NPs with high MPLA and CpG density, however, triggered equivalent IFN-β and IL-12p70 responses to MPs with high CpG, but low MPLA density.

The equivalent responses to MPs and NPs with matched CpG density indicates that the property of CpG density plays a dominant role in driving IFN-β and IL-12p70 responses. Notably, MPLA-CpG-Dual PLPs also showed CpG density-dependent IL27 response similar to IL12p70, suggesting activation and secretion of multiple cytokines from IL12 cytokine families. IL27 is an important cytokine that has important roles in vaccine-induced T cell response - in terms of magnitude and memory response (*39*). We also saw strong synergy for IL-10 production, similar to IL-12 and IFN-b, and although MPs outperformed NPs for CpG production even when densities were matched, the synergistic response was driven by adjuvant density rather than particle size. Interestingly, for IL-6 and TNF-alpha responses, we saw that particle size had a stronger effect than adjuvant density. There was no synergy in dual-delivery, and for both cytokines, MPs were much more effective for CpG single-adjuvant delivery than NPs, regardless of CpG density. This difference was also similar for dual-adjuvant delivery. Furthermore, MPLA-alone induced significant IL-6 and TNF expression, unlike what we saw for IFN-b and IL-12; and for lower density of MPLA delivery as a single adjuvant, NPs outperformed MPs in IL-6 production. Taken together, these results further underscore the critical role of the biophysical properties of adjuvant presentation to APCs. Specifically, particle size and adjuvant density, and provides an additional way to fine-tune innate immunity induced by various TLR ligands, either as a mono-adjuvant or as combination adjuvants.

For mechanistic understanding of the large synergy seen in the Type-I interferon and IL-12p70 responses to dual-presentation of MPLA and CpG, we first evaluated if the adaptor proteins that are commonly known to play major roles in TLR4 and TLR9 individual signaling are playing a critical role in developing the synergistic immune response in BM-APCs. TLR4 is present in both cell-surface membranes and in the endosomal membranes. On the cell-surface, TLR4 activation by lipopolysaccharides or MPLA leads to the recruitment of the adaptor protein MyD88 and its associated myddosome proteins(*40*). In the endosome, TLR4 activation results in the recruitment of the TRIF adaptor protein, which signals through receptor-interacting serine/threonine-protein kinase 1 (RIPK1) to activate TRAF6. Both MyD88 and TRIF are critical to maximize TLR4-mediated dendritic cell maturation (*41*). TLR4-MyD88 interactions on the plasma membrane result in early innate immune activation, while TLR4-TRIF interactions in the endosome result in delayed innate immune activation.(*42*) In this study, we observed that MyD88 is the primary adaptor protein for TLR9-driven signaling by CpG, whereas TRIF and TLR4 are both required for IFN-β and IL-12p70 synergistic responses to MPLA-CpG dual engagement. This indicates that PLPs, which are rapidly ingested by APCs, may activate endosomal TLR signaling and the dual presence of TLR4 and TLR9 in the endosomes drives synergistic behavior. It should also be noted that cell-surface TLR4 signaling is expected to low at early time points because the MPLA is encapsulated inside the particles, while acidic environments (as in endo/lysosomes) can accelerate PLGA degradation (*43*) and release the MPLA more rapidly. In TLR9^mut^ BM-APCs, we observed synergistic responses from MPLA and CpG treatment that were lower than observed from wild-type BM-APCs. The TLR9^M7NBtlr/MmJax^ (TLR9^mut^) strain has a chemically induced single point mutation that prevents TNF-α production after CpG stimulation in macrophages (*44*). Given that TLR9 is mutated and not completely knocked down in this strain, it is likely that high doses of CpG can still trigger low amounts of TLR activation.

We showed that interferon regulatory factors downstream play a role in MPLA and CpG-mediated signaling. Knockdown of IRF3 in BM-APCs amplified the IFN-β and IL-12p70 responses to MPLA and CpG. IRF3 is known to inhibit the binding of IRF5 to IL-12 promoters, so IRF3 knockdown may upregulate the magnitude of IRF5 signaling, causing higher IFN-β and IL-12p70 response in IRF3^−/−^ BM-APCs (*45*). IRF7 knockdown did not affect the magnitude of the IFN-β response, but approximately halved the magnitude of the IL-12p70 response. In the IRF7^−/−^ BM-APCs, synergistic responses were still observed for both IFN-β and IL-12p70. Finally, and most importantly, IFN-β and IL-12p70 responses to both individual and combination adjuvants were ablated in IRF5^−/−^ BM-APCs. IRF5 knockdown has been shown to downregulate CpG-mediated pro-inflammatory cytokine production in haematopoietic cells (*46*). We also observed that knockdown of IFNAR results in an increase in levels of secreted IFN-β and IL-12p70. IFNAR typically internalizes free Type I interferons to perpetuate downstream signaling, so the absence of IFNAR would result in accumulation of extracellular IFN-β in the closed *in vitro* system where our experiments were conducted.

An in-depth analysis of IRF5 signaling revealed that synergistic responses to TLR4 and TLR9 activation are associated with prolonged signaling kinetics. After treatment with PLPs, we evaluated IRF5 phosphorylation, an essential event for nuclear translocation and subsequent transcription for Type I interferons and proinflammatory cytokines (e.g., IL-12p70). MPLA-MP upregulated levels of phosphorylated IRF5 in the window of 30 minutes to 1 hour after treatment of BM-APCs. Phosphorylation of IRF5 was delayed in response to MP-CpG; heightened levels of phosphorylated IRF5 were not observed until 24 hours after treatment. Strikingly, MPs with both MPLA and CpG upregulated levels of phosphorylated IRF5 from 30 minutes to 4 hours. This indicates that adjuvants with different signaling latency can be combined to form a system that drives a longer-lasting immune response. We discovered that prolonged IRF5 signaling was associated with a higher rate of IRF5 nuclear translocation. Analysis of fractionated lysates showed that nuclear IRF5 levels were significantly higher after treatment with MPLA and CpG than with either adjuvant alone. The time at which peak translocation occurs, 4 hours, is in agreement with studies evaluating NOD2 stimulation (*47*). Taken together, these data suggest that both prolonged phospho-IRF5 expression and increased amount of nuclear translocation are likely causes for the cytokine synergy observed.

Further, we studied the signaling kinetics of TRAF6 as it bridges TLR4 and TLR9 activation upstream with IRF5 signaling downstream. We found TRAF6 followed similar patterns (kinetics and magnitude of signaling) as IRF5 signaling for MPs with MPLA, CpG and dual adjuvants. MPLA stimulated BM-APCs upregulated at 30 minutes to 1 hour, while treatment with CpG peaked at 4 hours. Dual MPLA and CpG lead to a prolonged expression from 30 minutes to 24 hours as seen in protein levels via western blot. Overall, as with IRF5 signaling, elevated and sustained TRAF6 signaling from dual MPLA and CpG adjuvants aided in the synergistic cytokine response.

Previously, using a combination of soluble TLR adjuvants, such as LPS and CpG or Poly IC and CpG, Oyang et al. reported a synergistic increase in IL12p40 cytokine in mouse peritoneal macrophages (PECs) via MyD88-TRIF-IRF5 pathways (*48*). Our results show that in mouse BM-APCs, which is a mixture of bone marrow derived DCs, macrophages and monocytes, a similar signaling axis is active, and provides synergistic enhancement of both Type I interferon (IFN-beta) and IL-12p70 when MPLA and CpG are co-presented using synthetic polymer-based PLPs. Importantly, in this study, we identified the critical role of signaling kinetics and adjuvant density as key mediators of the synergy. These results can have significant implications in vaccine design given the relevance of DCs, monocytes and macrophages in eliciting vaccine responses and the potential of controlling synergistic responses by manipulating the biophysical properties of the adjuvant presentation.

Investigating innate immune responses for pathogen-like co-presentation of multiple PAMPs is an essential step in elucidating key components that modulate the host immune response against pathogens and allow us to develop better vaccines. Here we explored the effect of biophysical properties (adjuvant density and size) of MPLA-CpG-Dual pathogen-like particles on synergistic cytokine responses in BM-APCs in our study. Our results indicate that the density of CpG on PLPs is the main driver governing the synergistic IFN and IL-12 response seen, while PLP size also plays an important role in the production of other cytokines. Furthermore, our examination of TLR4 and TLR9 signalling pathways individually and in combination via adjuvant co-loaded PLPs yielded critical insights on how key intermediary proteins drive synergistic cytokine responses. Through gene knockout studies, and protein and cytokine analyses down the signalling cascade, we found that the adaptor protein TRIF is required for any synergy, and downstream sustained and elevated TRAF6 signalling and IRF5 phosphorylation are the key drivers for the magnitude of the synergistic cytokine response for MPLA and CpG co-presentation on PLPs. Overall, we have identified the principal signaling components (TRIF-TRAF6-IRF5) that drives synergistic cytokine response for PLP-presented MPLA and CpG adjuvant combination in BM-APCs.

## Materials and methods

### Study design

In this study, we co-presented two clinically-relevant TLR-based adjuvants, MPLA (TLR4 agonist) and CpG (TLR9 agonist) on our pathogen-like particles (PLPs) to GMCSF-differentiated (7-day culture) bone marrow-derived antigen presenting cells (BM-APCs) to understand the effect of biophysical properties of PLPs on the synergistic innate immune response of MPLA-CpG dual adjuvants. Further, our overarching goal was to identify key molecular mediators and mechanistically understand their roles for the synergistic innate immune response of MPLA-CpG-Dual PLPs in BM-APCs. We synthesized polymeric micro and nanoparticles and co-loaded with MPLA (encapsulation) and CpG (surface loading) at different densities (high and low) and treated BM-APCs with the same MPLA to CpG dose ratio (1:10) across various PLP formulations. Using BM-APCs from wild type mice, we examined the effect of biophysical properties on PLPs with MPLA and/or CpG on multiple pro-inflammatory cytokine responses, including IL-12p70 and IFN-beta. Further, we used BM-APCs from different TLR4 and TLR9 signaling pathway-specific gene knock-out mice to identify key molecular mediators of synergistic IL-12p70 and IFN-beta response induced by MPLA-CpG-Dual PLPs. Finally, we studied the signaling kinetics in wild type BM-APCs using western blot techniques to understand the molecular mechanism for the synergistic innate immune response for MPLA-CpG co-presentation on PLPs.

### Animals

All animal experiments were conducted in accordance to approved IACUC protocols by Georgia Institute of Technology and Emory University. Female, 8-12 weeks old C57/Bl6 mice (Jackson Labs, Bar Harbor, ME) were used for all wild-type studies. IRF3^−/−^, IRF5^−/−^, IRF7^−/−^, MyD88^−/−^, TRIF^−/−^, and IFNAR^−/−^ mice were bred and housed at Emory University. TLR4^lps-del/JthJ^ and TLR9^M7Btlr/MmJax^ mice (henceforth referred to as TLR4^mut^ and TLR9^mut^ mice; see supplemental methods for detailed description for these mice) were also purchased from Jackson Laboratory and subsequently housed at Emory University.

### Mouse BM-APC culture

Bone marrow was harvested from the tibias and fibulas of C57/Bl6 mice (6-10 weeks, Jackson Labs, Bar Harbor, ME). The bone marrow cells were processed through a 40 μm cell strainer, treated with RBC lysis buffer, and seeded in Petri dishes at a concentration of 1,000,000 cells/ml (total 20 million cells/Petri dish). BM-APCs were differentiated from bone marrow-derived cells through culture in RPMI media (Invitrogen) with 10% characterized fetal bovine serum (HyClone, Logan< UT), 1% penicillin-streptomycin, 2 mM glutamine, 1x β-mercapethanol, 1 mM pyruvate, and 20 ng/mL mouse recombinant GM-CSF (Peprotech, Rocky Hill, NJ). 10 mL media (50% of initial volume in Petri dish) was replaced with fresh media supplemented with GMCSF (40 ng/mL) on days 2 and 4. On day 6, 10 ml GMCSF-supplemented (at 60 ng/mL) media was added without replacing any media. On day 7, BM-APCs were harvested and replated in fresh culture media supplemented with GMCSF at 20 ng/mL for further experiment using PLP adjuvants.

### Pathogen-like particle (PLP) synthesis and characterization

TLR9 adjuvant ODN1826 was purchased from InvivoGen (San Diego, CA). Monophosphoryl Lipid A (Salmonella Minnesota R595) was purchased from Avanti Polar Lipids (Alabaster, AL; Cat#699200P). PLGA microparticles (MPs) and nanoparticles (NPs) were synthesized using a double emulsion method as reported previously by us.(*33, 49*) Resomer ® RG 502 H Poly(D-Lactide-Co-Glycolide) (Sigma-Aldrich, St. Louis, MO) was dissolved in dichloromethane and water was added and homogenized to form a primary emulsion. For MPLA formulations, MPLA was added to the dichloromethane before emulsification. MPs were formed through homogenization at 10,000 rpm for 2 minutes. The secondary emulsion was formed by adding the primary emulsion to 1% polyvinyl alcohol (87-89% hydrolyzed, Sigma Aldrich) solution and homogenizing at 10,000 rpm for 2 minutes. NPs were formed through sonication at 65% power for 2 minutes. The secondary emulsion was formed by adding the primary emulsion to a solution of 5% polyvinyl alcohol (87-89% hydrolyzed, Sigma Aldrich) and sonicating at 65% power for 5 minutes. Dichloromethane was removed through rotary evaporation for 3 hours. PLGA MPs were pelleted by centrifugation for 20 minutes at 3000 g and washed with DI water two times. PLGA NPs were pelleted by ultracentrifugation at 22,000 g for 20 minutes and washed with DI water 2 times. Both MPs and NPs were surface modified with branched polyethylenimine (Polysciences, MW=70,000, Warrington, PA) through reaction via EDC and sulfo-NHS with a non-toxic level of PEI conjugated to PLGA particles (~6.5 µg PEI/mg PLGA) as shown in previous works (*33, 49*). PEI-modified MPs were pelleted by centrifugation for 20 minutes at 3000 g, washed with 1 M NaCl solution 2 times, and washed with DI water once. PEI-modified NPs were pelleted by ultracentrifugation for 20 minutes at 22,000 g, washed with 1 M NaCl solution 2 times, and washed with DI water once. Both PEI-modified particle formulations were flash frozen in liquid nitrogen and lyophilized for 48 hours and stored at −20°C. Measurements of MP or NP size and zeta potential were performed in 1 mM KCl solution with a Malvern Zetasizer. Loading of CpG was quantified using the Nucleic Acid Quantification module of the Gen5 software on a BIOTEK Synergy HT Plate Reader.

### *In vitro* activation of mouse BM-APCs with PLP-adjuvant formulations

On day seven of culture, BM-APCs were plated at a density of 300,000 cells/well in 96-well plates and allowed to settle for two hours before the addition of PLP adjuvants. Cell density to particle ratio was preserved and extrapolated to encompass 6-well plates and 10 cm petri dishes for larger experiments. After treatment with PLP adjuvants, supernatants were harvested at 0.5, 1, 2, 4, 6, 12, and 24 hours. ELISA or Luminex (Bio-Techne, Minneapolis, MN) was used to measure cytokine concentrations after cell activation (IFN-β, IL-12p70, IL-6, TNF-α, IL-10). BM-APCs were lysed using Cell Lysis Buffer (Cell Signaling Technologies, Danvers, MA, Catalog # 9803S) supplemented with PMSF (Cell Signaling Technologies, Danvers, MA, Catalog # 8553S) to collect proteins for Western blots. For Phos-tag gels, BM-APCs were lysed using an EDTA free lysis buffer of 50mM Tris-HCl (pH 7.4), 150mM NaCl, 1% NP-40, 0.5% Sodium deoxycholate, 0.1 % SDS supplemented with complete protease inhibitor (Roche, Cat#11836170001) and PhosSTOP phosphatase inhibitor cocktail tablets (Roche, Cat#04906837001)(*50*). Cell fractionation kit (Cell Signaling Technologies, Danvers, MA, Catalog # 9038S) supplemented with PMSF and Protease inhibitor cocktail (Cell Signaling Technologies, Danvers, MA, Catalog, # 8553S; # 5871S) was used to separate nuclear from cytoplasmic proteins. Directions were followed as provided by the manufacturer.

### Western blots

Mouse BM-APCs were lysed in 1X Cell Lysis buffer (Cell signaling technologies, Catalog #9803) or EDTA free lysis buffer (*50*) or cell fractionation kit (CST) according to western blot; SDS-PAGE, Phos-tag or Nuclear fractionation; respectively. Appropriate volume of lamellae buffer was added to the lysate (2X for SDS-PAGE and Phos-tag samples and 6X for cell fractionation samples). Samples were mechanically homogenized by French press method using 28-gauge insulin syringe before boiling at 96 degrees for 10 minutes and cooled on ice. Equal lysate concentrations were then loaded onto a Bio-Rad mini PROTEAN TGX gel or 50 µM Phos-tag acrylamide gel and electrophoresis on either Bio-Rad gel tank or Life Technologies Mini gel tank, respectively. After gel electrophoresis, gel was transferred either by Bio-Rad Trans-Blot turbo or wet tank transfer at room temperature onto a PVDF membrane following the protocol provided for each gel type (SDS-PAGE or Phos-tag). Transferred PVDF was blocked with 5 % Bio-Rad blocking buffer milk (Catalog # 170-6404) in TBST for 3 hours, rocking at 4 degrees. Then subsequent primary antibody incubation was done overnight at 4 degrees (all primary antibodies were diluted in 1% blocking milk or for Phos-tag with MBL Max blot solution 1 or Takara Western blot immune booster). Proteins of interest were detected with Anti-Rabbit IgG HRP (Cell signaling technologies) and visualized with Bio-Rad Clarity Max according to manufacturing protocol.

Antibodies purchased from Abcam, IRF5 Chip grade (catalog#ab21689, final dilution 1/3000), Cell signaling technologies: IRF5 (Catalog#4950S, final dilution 1/1000), IRF3 (Catalog#4302S, final dilution 1/1000), Thermo/Invitrogen: Phosphorylated IRF3 (Ser 396) (Catalog#MA5-14947, 1/1000 dilution) and TRAF6 rabbit monoclonal (Catalog#702286, 1/1000 dilution). Secondary antibodies were from Cell signaling technologies: Anti-Rabbit-HRP-labeled (catalog#7074P, 1/10000 dilution).

Loading controls were visualized by either Invitrogen Gapdh (GA1R) mouse monoclonal (Catalog#MA5-15738, 1/5000 dilution), Monoclonal Beta-Actin-Sigma (Catalog#A5316, 1/5000 dilution), Cell Signaling Technologies-Rabbit monoclonal of Vimentin (D21H3)-(Catalog#5741, 1/1000 dilution), Histone (D1H3) (Catalog#4499, 1/1000 dilution), Vinculin (4650, 1/1000 dilution).

### Statistical analysis

All statistical analyses were performed using GraphPad Prism 8. Data are presented in bar graphs as mean ± standard deviation (SD). To assess the statistical significance of the difference between three or more normal datasets, a one-way ANOVA was performed. Multiple comparisons were evaluated using Tukey’s test and P values less than 0.05 were considered significant between two groups.

## Acknowledgements

We acknowledge the assistance of Cedrick Young and Jia Yao in maintenance of the breeding colony for knockout mice used in these studies, M. Cole Keenum for assistance with particle synthesis and characterization, and Neha Narang and Gabriela Vogel for technical assistance with studies evaluating immune response kinetics. We acknowledge funding support from NIH grant U01-AI124270-02 and the Robert A. Milton Chaired Professorship to Krishnendu Roy.

## Author Contributions

PP, RT, KR conceptualized studies. PP, RT, NJ, ELB, AA, BP performed the experiments. PP, RT, NJ, KR wrote the manuscript. PJS, DMS and KR provided resources, guidance and designed experiments.

## Competing Interests

The authors declare no competing interests.

## Notes

### Competing Interest Statement

The authors have declared no competing interest.

