## Supplemental Information for "TRAF6-IRF5 kinetics, TRIF, and biophysical factors drive synergistic innate responses to particle-mediated MPLA-CpG co-presentation"

#### Supplemental Method

***Tlr4**^lps-del^**/JthJ*** **mouse from Jackson Laboratory (**https://www.jax.org/strain/007227**):** The stimulation of Toll-like receptor 4 (TLR4) by lipopolysaccharide (LPS) induces the release of proinflammatory cytokines that activate immune responses. The $Tlr4^{Lps\text{-}del}$ spontaneous mutation corresponds to a 74723 bp deletion that completely removes the *Tlr4* coding sequence. No mRNA or protein is expressed. Homozygous mutants exhibit a defective response to LPS stimulation. The functionally similar $Tlr4^{Lps\text{-}d}$ mutation found in C3H/HeJ mice (#000659) is a point mutation that causes an amino acid substitution.

**$TLR9^{M7Btlr/MmJax}$ mouse from Jackson Laboratory** (https://www.jax.org/strain/014534): Mice that are homozygous for this mutation are viable, fertile, normal in size and do not display any gross physical or behavioral abnormalities. In response to stimulation with oligodeoxynucleotides containing CpG motifs, macrophages do not produce tumor necrosis factor (TNF) alpha. This mutant mouse strain may be useful in studies of the role of toll like receptor 9 (TLR9) in the immune system.

### Supplemental Table

#### Table S1. PLP formulations for MPLA and CpG

| TLR adjuvants and Loading methods | **MP** formulations | **NP** formulations (Low density) | **NP** formulations (High density) |
|---|---|---|---|
| **MPLA** (M)<br><br>*Encapsulation* | MLo MP<br><br>*Density: 1.34E-06 mg/m3*<br>*Loading: 1 ug MPLA /mg MP* | MLo NP<br><br>*Density: 1.34E-06 mg/m3*<br>*Loading: 1 ug MPLA /mg NP* | MHi NP<br><br>*Density: 8.04E-06 mg/m3*<br>*Loading: 6 ug MPLA /mg NP* |
| **CpG** (C)<br><br>*Surface loading* | CHi MP<br><br>*Density: 3.35 mg/m2*<br>*Loading: 10 ug CpG /mg MP* | CLo NP<br><br>*Density: 0.056 mg/m2*<br>*Loading: 10 ug CpG /mg NP* | CHi NP<br><br>*Density: 3.35 mg/m2*<br>*Loading: 60 ug CpG /mg NP* |
| **MPLA** (M) + **CpG** (C)<br><br>*Encapsulation - MPLA*<br>*Surface loading - CpG* | MLo-CHi-Dual MP<br><br>*Density and loading - same as MLo-MP and CHi-MP* | MLo-CLo-Dual NP<br><br>*Density and loading - same as MLo-NP and CLo-NP* | MHi-CHi-Dual NP<br><br>*Density and loading - same as MHi-NP and CHi-NP* |
| *Dose of adjuvants- MPLA at 50 ng/ml and CpG at 500 ng/ml concentration*<br>*Dose ratio - MPLA:CpG is 1:10* | | | |
| *Note, it was not possible to prepare MHi-CHi-Dual MP or MLo-CLo-Dual MP formulations that require co-loading of MPLA (by encapsulation) and CpG (by surface loading) on MPs at 1:10 weight ratio between MPLA and CpG (to maintain same adjuvant doses across MP and NP formulations) with 100% loading efficiency for both adjuvants to match adjuvant densities on the corresponding NP formulations. For our formulations, approximately 6-fold higher surface area for NPs (avg size – 250 nm) compared with MPs (avg size - 1.5 um) enables 6-fold higher surface loading of CpG on NPs with 100% efficiency. Typically, our MPs can surface load up to a maximum of 10 ug CpG/mg particle with 100% efficiency whereas NPs can load up to 60 ug CpG/mg particle with 100% efficiency due to 6X higher surface area. 100% loading efficiency of adjuvants is important to maintain the dose ratio between MPLA and CpG across experiments.* | | | |

**Table S2. Size, zeta potential and loading levels for PLP formulations**

| **PLP formulations** | **Size (nm)** (before PEI modification) | **Zeta (mV)** (after PEI modification) | **Loading levels for MPLA and/or CpG** |
|---|---|---|---|
| MLo MP | *1353.3 ± 154.5 (n=3)* | *+28.9 ± 1.6 (n=3)* | *1 ug MPLA /mg MP* |
| CHi MP | *1114.6 ± 336.2 (n=3)* | *+29.4 ± 3.2 (n=3)* | *10 ug CpG/mg MP* |
| MLo-Chi-Dual MP | *1353.3 ± 154.5 (n=3)* | *+28.9 ± 1.6 (n=3)* | *1 ug MPLA and 10 ug CpG /mg MP* |
| MLo NP | *240.3 ± 4.8 (n=4)* | *+28.5 ± 2.1 (n=4)* | *1 ug MPLA /mg NP* |
| CLo NP | *248.2 ± 31.2 (n=4)* | *+29.1 ± 0.8 (n=4)* | *10 ug CpG/mg NP* |
| MLo-CLo-Dual NP | *240.3 ± 4.8 (n=4)* | *+28.5 ± 2.1 (n=4)* | *1 ug MPLA and 10 ug CpG /mg NP* |
| MHi NP | *238.8± 35.3 (n=2)* | *+26.3 ± 0.4 (n=2)* | *6 ug MPLA /mg NP* |
| CHi NP | *248.2 ± 31.2 (n=4)* | *+29.1 ± 0.8 (n=4)* | *60 ug CpG/mg NP* |
| MHi-CHi-Dual NP | *238.8± 35.3 (n=2)* | *+26.3 ± 0.4 (n=2)* | *6 ug MPLA and 60 ug CpG /mg NP* |

### Supplemental Figures

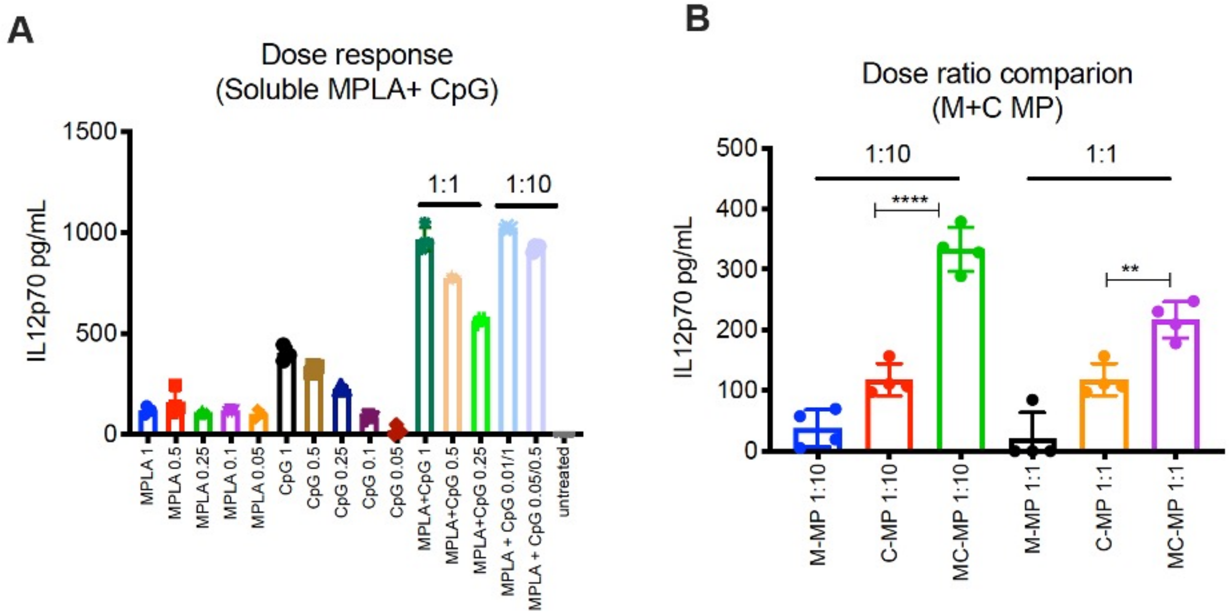


**Figure S1.** Effect of various doses and ratios of soluble and MP formulations of MPLA and CpG on IL-12p70 secretion by BM-APCs. Data represent Mean ± SD. **P < 0.01, ****P<0.0001, one-way ANOVA with Tukey's multiple comparison test.

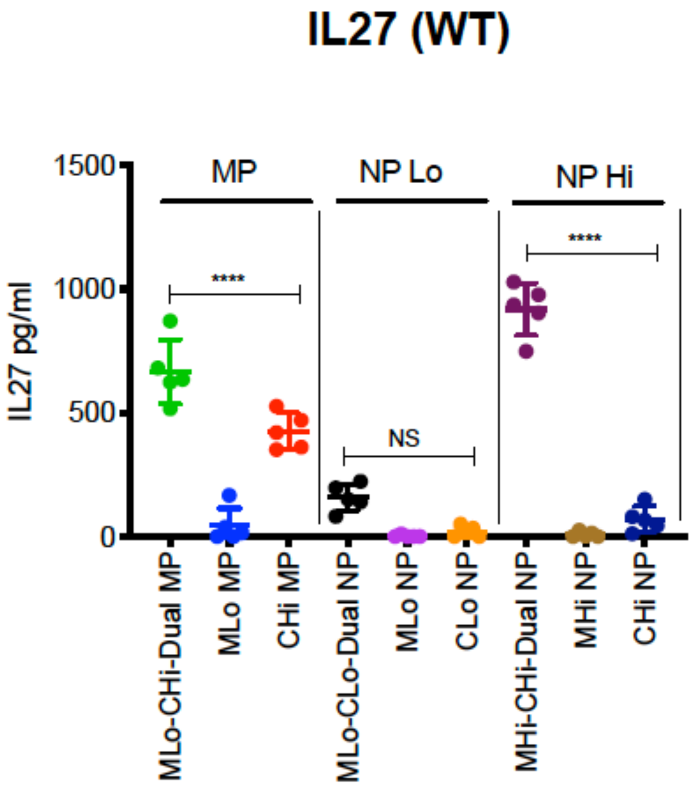


**Figure S2.** Synergistic IL27 response from BM-APCs induced by pathogen-like particles (PLPs) with MPLA and CpG depend on CpG adjuvant density. Center lines designate the mean value and error bars represent SD. ****P<0.0001, NS- not significant; one-way ANOVA with Tukey's multiple comparison test.

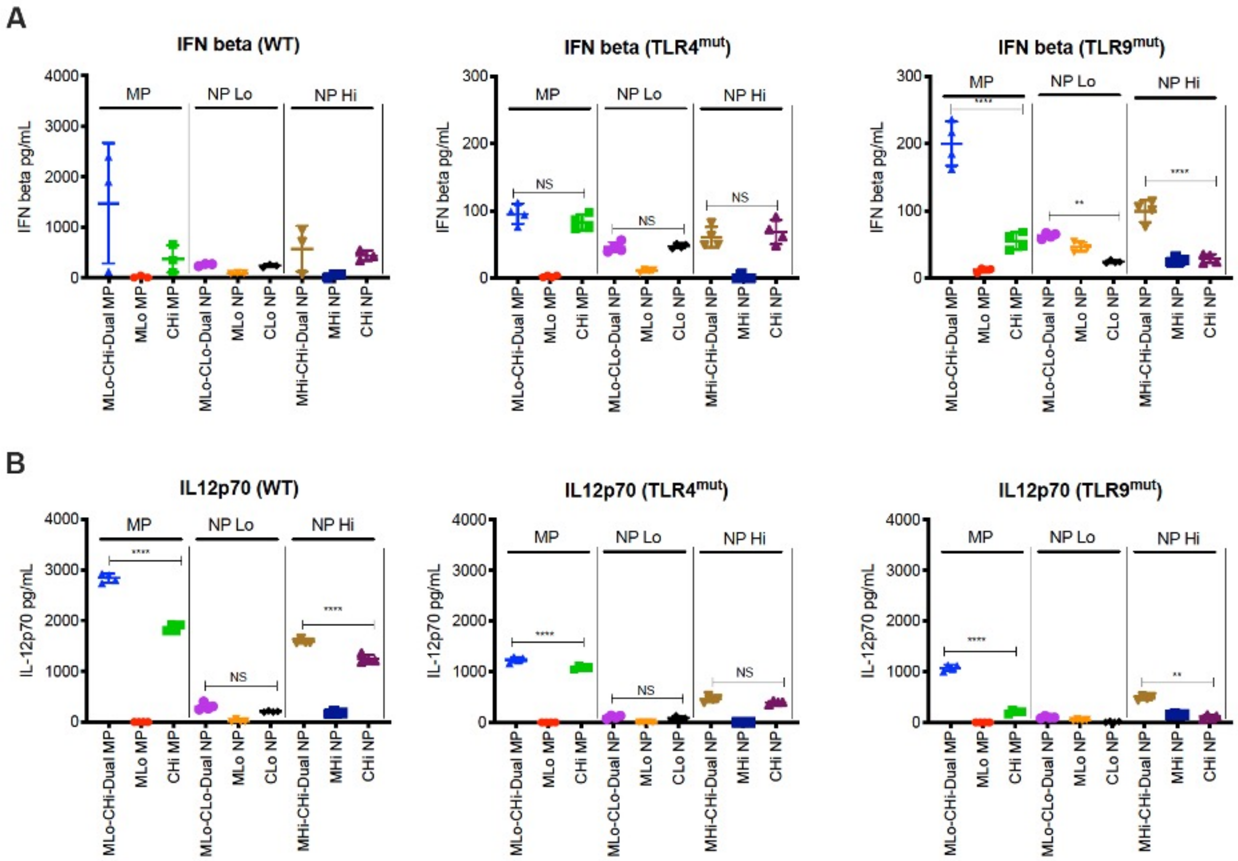


**Figure S3.** A) IFN-β and B) IL-12p70 secretion from BM-APC's after PLP treatment in wild-type, TLR4$^{mut}$, and TLR9$^{mut}$ BM-APCs. Center lines designate the mean value and error bars represent SD. **P < 0.01, ****P<0.0001, NS- not significant; one-way ANOVA with Tukey's multiple comparison test.

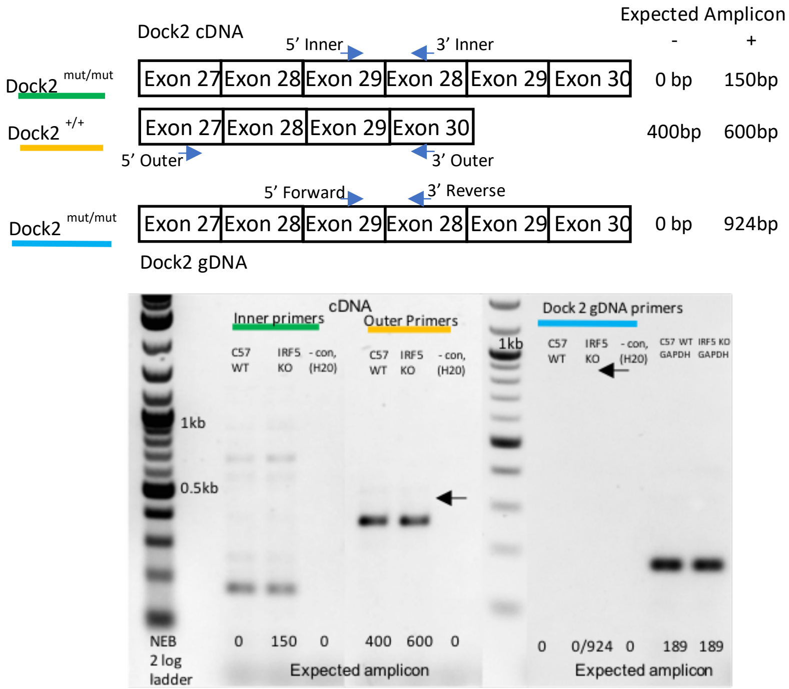


**Figure S4:** Possibility of DOCK2 mutation in our IRF5$^{-/-}$ mice colony. Amplicon was visualized via 1.5% agarose gel. **Lane 1**: NEB 2 Log ladder. **Lane 3 and 4**: Inner primers (spanning Exon 29 and 28) on cDNA flanking the DOCK2 mutation. An amplicon is visualized at 150bp in both wild type C57 and IRF5$^{-/-}$ samples. If there was DOCK2 mutation, then amplicon would only be seen in the IRF5$^{-/-}$. To reassess this amplicon, new gDNA primers were used to reassess this mutation (Lane 10 and 11). **Lane 6 and 7:** Outer primers (spanning Exon 27 and 30) on cDNA. Amplicon at 400 bp in conjunction with no amplicon at 600bp indicates no DOCK2 mutation in our IRF5$^{-/-}$ colony. **Lane 9:** NEB 2 Log ladder. **Lane 10 and 11:** new genotyping primers with gDNA for DOCK2 mutation in wild type and IRF5$^{-/-}$ mice. Lack of 924bp amplicon indicates no DOCK2 mutation in our IRF5$^{-/-}$ colony. This was reconfirmed with Purtha et al. (*1*). All primers and schematic are adapted from Purtha et al. (*1*).

cDNA Primers:

Outer forward 27–30, 5′- GGATGCGGCCTTCACTTA-3′; and Outer reverse 27–30, 5′- TCCACAGCTGGAACTC- AAAG-3′.

Dock2 Mutation forward 29–28, 5′-CAAGGACCTCATTGGGAAGAA-3′; and Dock2 mutation reverse 29–28, 5′-CTGAGCTGGTCTGGAAGGTCT-3′

gDNA Primers- Dock 2 mutation forward: TCACTGCCCCTTAATGATGTC, Dock2 mutation reverse: TTGCCTTTGACACACCGTAG

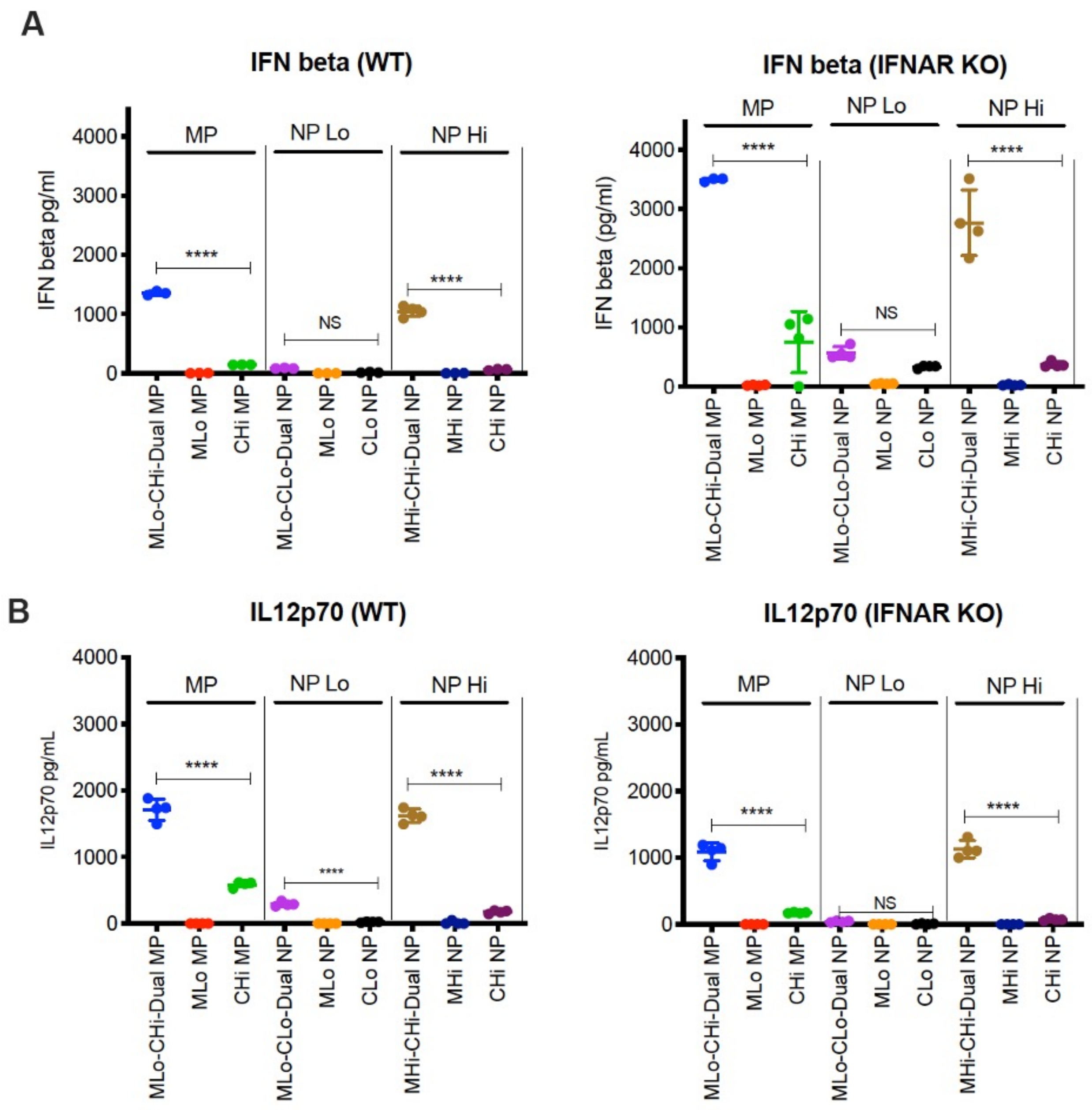


**Figure S5.** IFN-β and IL-12p70 secretion from BM-APCs after PLP treatment in wild-type and IFNAR knockout BM-APCs. Center lines designate the mean value and error bars represent SD. ****P<0.0001, NS- not significant; one-way ANOVA with Tukey's multiple comparison test.

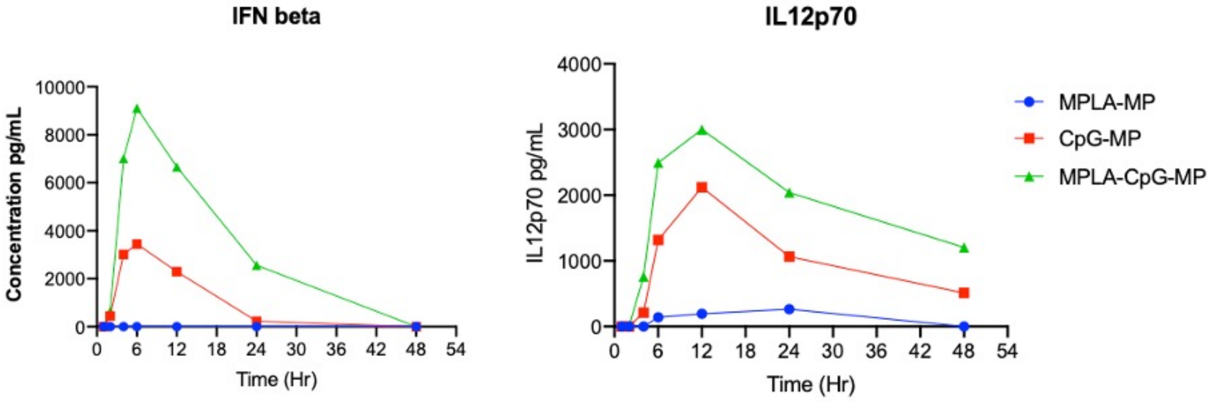


**Figure S6.** Kinetics of IFN beta and IL-12p70 production by BM-APCs treated with PLPs. BM-APCs from this experiment was used for Nuclear fractionation studies and the results are shown in **Figure 4**.
